# Evaluation and primer for using DeepLabCut to track small animals in large spaces

**DOI:** 10.64898/2026.10.15.759451

**Authors:** Lata Kalra, Susan Balenger, Norman Lee, Swastik P. Padhy, Mingzi Xu

**Author notes:** Correspondence: Lata Kalra, Department of Ecology, Evolution, and Behavior, University of Minnesota – Twin Cities, St. Paul, MN, U.S.A.

## Abstract

DeepLabCut (DLC), a machine learning-based pose estimation tool, has dramatically increased the throughput of behavioral tracking. Yet its implication remains limited by lack of validation data from diverse scenarios and users’ unfamiliarity with the machine learning workflow. Here, we provide a practical primer on DLC terminology and workflow and performance evaluation of DLC in a novel scenario where the target is much smaller than those in previous validation data. We assessed model performance in a total of 36 model configurations across three training proportions, six training iterations, and two recording resolutions. Performance was evaluated on training convergence, prediction error, prediction certainty, and generalizability. Additionally, we selected an optimal model based on a balance between model performance and training time and applied it to 20 recordings at each resolution. In the test frames, 15 and 6 out of 18 model configurations met all performance criteria under high-and low-resolution, respectively. Of these, DLC was able to achieve low prediction error, high prediction confidence, and low train-test loss. Prediction error and confidence varied with training proportion, iteration, and resolution. Inference using the optimal models achieved similar high confidence predictions, with low-likelihood predictions predominantly associated with visual obstruction, low light, and reduced contrast between the subject and the background. Notably, the optimal models for high-and low-resolutions performed similarly despite a multi-fold difference in the cost of the recording devices, providing validation data for DLC in low-budget research, teaching and outreach scenarios.

## Introduction

Behavioral tracking is a pivotal component of research and teaching in multiple areas of biology, such as tracking foraging trajectories (Haalck et al., 2026), quantifying movement responses to courtship songs (Gerhardt and Huber, 2002), and monitoring locomotor outputs of neural circuits (Bogachev et al., 2023). Many animal behaviors are complex and fast, and hence, tracking and extracting information from them in real time can be challenging. Tracking behaviors from video recordings is frequently used to solve this issue. Traditionally, videos are analyzed manually. While this approach allows a high degree of user-customization and can be feasible in field conditions due to simple facility requirements, it can be time-consuming, error-prone, and may introduce both within-and between-observer bias (Levitis et al., 2009). Recently, machine learning (ML)-based programs, especially open-source programs such as DeepLabCut (DLC; Mathis et al., 2018; Nath et al., 2019), are becoming an increasingly popular tool for behavioral tracking. Compared to manual tracking, tracking using ML-based programs is much time-and labor-efficient while achieving error rates comparable to human observers (Correia et al., 2024; Hayakawa et al., 2024; Mathis et al., 2018; Nath et al., 2019; Sturman et al., 2020).

ML-based behavioral tracking programs can be trained to detect gestural and movement patterns of interest within images or videos using Convolutional Neural Network (CNN; Box 1B), a category of artificial neural network (ANN) specialized for image recognition (Deng, 2014). The markerless pose estimation workflow (Box 1A, 1B) enables users to manually label focal body parts as ground-truth (Box 1B) keypoints on a small subset of the video frames (Fig. 1A,B). These labelled frames can then be split into “train” and “test” frames (Box 1B; Fig. 1C; Greener et al., 2022; Nath et al., 2019). Training in ANNs is iterative, which allows learning to occur. This process usually involves training the algorithm to track focal body parts in the “train” frames and evaluating the model performances in the “test” frames, and subsequently, adjusting various training parameters (Box 1B, Box 2) until satisfactory performance is achieved (Fig. 1D,E). Once trained, the learned model can be used to detect the same patterns in new videos (Fig. 1F), offering a major advantage over manual tracking in terms of time-efficiency, error rate, and biases in data extraction (see Box 1A for description of entire workflow).

**Figure 1:**
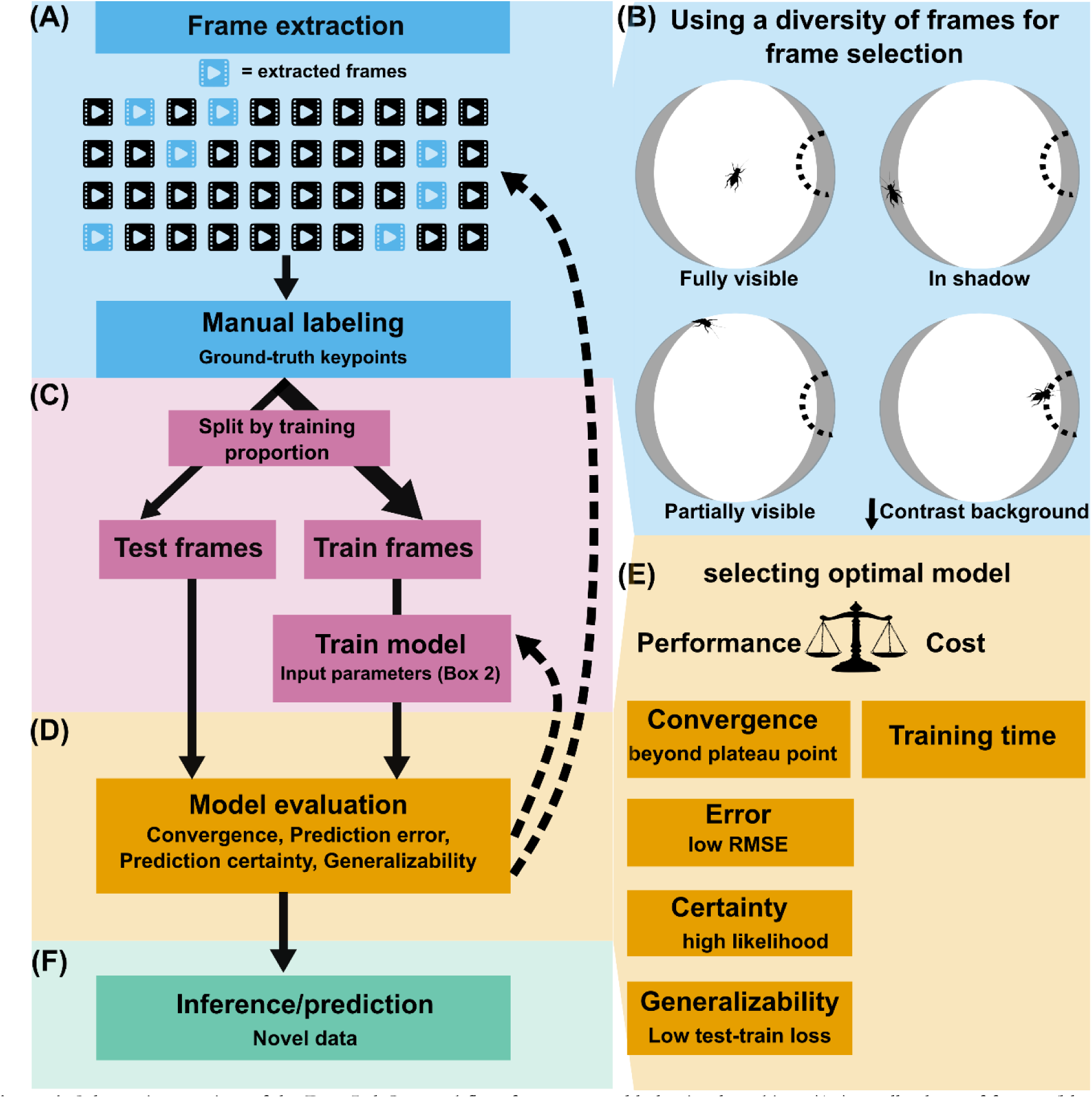
Schematic overview of the DeepLabCut workflow for automated behavioral tracking. A) A small subset of frames (blue, 100-200 total) is extracted from the full video dataset and manually labeled to create ground-truth keypoints. B) It is advisable to include a diverse set of frames, including those where subject-background contrast is low or where subject is partially visbile or in poor light conditions. C) Labeled frames are then split into training and test subsets based on a user-defined training proportion, for example, 95%, 80% and 50%. The model is trained on the training frames while test frames are withheld at this stage. D) At the evaluation stage, model predictions for both train and the held-out test frames are compared against ground-truth labels and models can be retrained with different paramters or with a different set of extracted frames (dashed curved arrows) until satisfactory performance is reached. E) An optimal model configuration can then be determined based on a balance between performance an computational cost. Metrics like achieving training convergence, low error and high certainty of predictions and high model generalizability can be used to evalaute performance while computational cost can be estimated using metrics like total training time. F) Once an optimal model is selected, the entire dataset of frames can then be analyzed in the inference stage.

Although ML-based automated behavioral tracking offers multiple benefits, several barriers limit its application in research and teaching. First, lack of familiarity with the machine learning workflow and terminology may present a steep learning curve to potential users with no prior exposure to machine learning. Second, published validation data on the performance of ML-based programs have limited application scope. To date, validation data are predominantly from scenarios wherein the focal object occupies a substantial portion of the video frame, for instance, a fly enclosed in a small arena or a zoomed-in forelimb of a mouse on a joystick (Arent et al., 2021; Brandt et al., 2021; Chanthongdee et al., 2024; Correia et al., 2024; Hardin and Schlupp, 2022; Hayakawa et al., 2024; Jernigan et al., 2023; Lauer et al., 2022; Mathis et al., 2018; Nath et al., 2019; Sturman et al., 2020a; Wiltshire et al., 2023). However, many ecologically relevant behaviors like mate searching, foraging, migration, and predation often require tracking the animals across spaces that are multifold larger than their body sizes. Data to evaluate ML-based tracking programs’ performance in such scenarios is lacking.

To overcome these two barriers, in this study, we provide a practical primer to demystify concepts, workflow, and terminology of ML-based behavioral tracking for biologists with an emphasis on DeepLabCut (DLC) and offer evaluation of DLC’s performance in tracking small animals in relatively large spaces using the mate-searching behavior of female crickets as an example. We evaluate DLC performance on both high-resolution videos from a 4K video camera and low-resolution videos from a low-cost webcam, representing equipment set-up in well-funded research labs and teaching labs limited in funding, respectively. For both recording resolutions, we investigated the effect of training proportion (Box 1B, proportion of labelled frames used for training) and training iterations (Box 1B) across four aspects of performance: convergence (Box 1B), prediction error, prediction certainty, and generalizability (Box 1B). In addition, we apply a strategy balancing performance and time cost to select optimal model for use of inference (Fig. 1E) and provide diagnosis of common erroneous scenarios in model inference.

## Primer on DeepLabCut

#### BOX 1A: DLC WORKFLOW

DeepLabCut (DLC) is a machine learning-based, open source, markerless pose estimation (Box 1B) software package adapted from the multi-person pose estimation tool DeeperCut (Lauer et al., 2022; Mathis et al., 2018; Nath et al., 2019). DLC enables researchers to train models on relatively small, custom datasets with high accuracy, making it a flexible tool for quantifying animal movements. The typical workflow (Fig. 1) for tracking using DLC includes four steps: (1) project setup and manual labeling, (2) model training using manually labeled frames, (3) model evaluation and refinement, and (4) applying the trained model to new videos (also referred to as, inference; see Box 1B).

(1) **Project setup and manual labeling:** users begin the workflow with project creation, which sets up a project folder and generates a configuration file (config.yaml; Box 1B). This configuration file contains input parameters the program uses in training, such as experiment name, video paths, and which body parts to track (see Box 2 for details on input parameters). After the project is set up, users can execute frame extraction (Box 1B) using methods such as uniform sampling and K-means clustering (Box 1B), and manually label focal body parts using DLC’s graphical user interface (GUI) on a small subset of frames. These manual labels are used both for training the model to recognize the focal body parts and as the ground-truth (Box 1B) keypoints in evaluating model performance in step (3); labels are saved as x and y coordinates (Box 1B). Frame extraction should consider good coverage of poses, orientations of the target body parts, and backgrounds to achieve better model generalizability (Fig. 1B). Once the extracted frames are labeled, they are split into test and train frames (Box 1B) based on the designated training proportion. Multiple shuffles (Box 2) of splitting the extracted frames can be implemented for training. Approximately 100-200 labeled frames are often sufficient for accurate tracking (Mathis et al., 2018; Nath et al., 2019).
(2) **Training of the model using manually labeled frames:** DLC uses convolutional neural network (CNN; Box 1B) to train the model. In plain language, training the model means the model parameters (also called network weights, Box 1B) are being adjusted such that predictions made by the model become increasingly accurate. CNNs used by DLC come pretrained with large scale image databases to recognize general visual features (e.g., edges, corners, shapes). The model is then fine-tuned using manually labeled keypoints from step (1). This fine-tuning step, termed transfer learning (Box 1B), allows the model to be adapted to a specific project. For example, after fine-tuning, DLC learns things such as appearance of the focal animal or body part in the video and its variations, relationships among body parts, and features that distinguish the target from the background in the project-specific videos. Training can be computationally intensive and is ideally done on a graphical processing unit (GPU; Box 1B) or cloud computing platforms (e.g. Google Colab). It can be paused, resumed, and progress can be monitored through cross-entropy loss curves (Box 1B). Training convergence indicates the point beyond which further training is likely to yield diminishing returns.
(3) **Model evaluation and refinement:** once a model is trained, DLC offers metrics to evaluate model performances. These data include: Root Mean Square Error (RMSE) for training and test frames, likelihood, cross-entropy loss, and test-train loss (see Box 1B for all terms). DLC also offers an option to visually inspect model predictions in videos and overlay predictions on test images. The workflow bifurcates after evaluation. If model performance is satisfactory, users can proceed to use the trained model to make predictions in new videos. If model performance is not satisfactory, users can adjust input parameters (e.g., training proportion or iterations), add or replace manual labels, and retrain the model. This process can be iterative until performance is satisfactory. (*Continued on next page*)
(4) **Applying the trained model to new videos:** once a satisfactory model is obtained, it can then be applied to new videos. The model output will include both CSV and HDF5 files (Box 1B) that contain x- and y-coordinates and likelihood values for each labeled body part in every frame. Successful application of DLC typically depends on video quality. Resolution should be sufficient to resolve tracked body parts of interest, and frame rate should match the temporal scale of the behavior. Even illumination and background that provides high contrast with the target can generally improve tracking performance. When combined with thoughtful project setup and iterative refinement of labeled frames, DLC provides a powerful, efficient, and generalizable platform for automated extraction of detailed, quantitative behavioral data.

#### BOX 1B: Glossary for DLC Workflow

(1) **Project setup and manual labeling:**

**Markerless pose estimation:** estimation of the positions of anatomical landmarks in a video, without physical markers, using computer vision models trained on labeled images.

**YAML file (config.yaml):** a human-readable configuration file used for users to specify information about project settings (e.g., file paths, body parts to track, training parameters; see Box 2). DLC uses information in this file to construct and train models.

**Frame extraction:** selecting a subset of frames from videos for manual labelling of the individual(s) or body part(s) of interest.

**Uniform sampling:** extracting frames at regular intervals from a video to avoid temporal bias in the training set. **K-means clustering:** a method DLC can use to group similar frames; selecting frames from different clusters increases diversity in frames selected for manual labeling.

**Ground truth:** the set of manually labeled keypoints that serve as the reference against which model predictions are compared.

**x, y coordinates:** pixel-space positions of each labeled body part in a frame

**Graphics Processing Unit (GPU):** parallel processor designed to handle large numbers of operations simultaneously. In machine learning, using a GPU can accelerate deep network training and inference, substantially reducing run time compared to a CPU.

**Train frames:** the subset of manually labeled frames used to train the model.

**Test frames:** the subset of manually labeled frames withheld from training. After training, test frames are used to evaluate the trained model’s predictions, which offers indications about how well the trained model may perform on new, unseen data.

(2) **Training of the model using manually labeled frames:**

**Convolutional neural networks (CNNs):** neural networks specialized for images; they learn hierarchical visual features via convolutional layers (see below) to detect body parts and poses.

**Convolutional layers:** layers within the CNN that extract different visual features. Early layers detect simple features such as edges, lines, corners, and orientations and deeper layers process complex patterns such as shapes, contours, and body parts.

**Network weights**: numerical parameters that encode network learning. The network weights are adjusted in each cycle of training to customize the network to a specific project such that prediction error can be reduced.

**Transfer learning:** reusing a network pre-trained on large image datasets and fine-tuning it with a set of small, task-specific and labeled frames to achieve high accuracy with fewer labels.

**Training iteration**: one cycle in which the CNN processes a batch of labeled frames, computes the prediction error, and adjusts the network weights via transfer learning to reduce errors. Model weights at each iteration can be saved as snapshots at regular intervals for evaluation of models at different stages of training.

**Training proportion:** the proportion of labeled frames that are designated for training the model. Commonly adopted training proportions include 50%, 80% and 95% (DLC default). For instance, for an 80% training proportion, 80% of manually labeled frames get assigned for training the model and the rest 20% frames are held from training and used for evaluating (testing) the model (see *TrainingFraction* Box 2).

(3) **Model evaluation and refinement:**

**Cross-entropy loss:** a unitless metric indicating how well the model’s predicted labels match the ground-truth labels during training. Lower values indicate better model performance. Cross-entropy loss typically follows an exponential decay pattern during training where a steep decrease across iterations indicates the model is still training, while a plateau indicates that training has stabilized (see Training convergence below).

*(continued on next page…)*

**Training convergence:** the point during training at which cross-entropy loss plateaus and additional iterations yield diminishing returns in model improvement. Convergence can be assessed by fitting a decay curve to the loss values and determining the iteration at which the curve enters the asymptotic plateau. Models trained beyond the convergence point are unlikely to improve substantially.

**Root Mean Square Error (RMSE):** the square root of the mean squared Euclidean distance (in the unit of pixels) between a model’s predicted and ground-truth keypoint locations. It is one of the fundamental metrics for evaluating model performance.

**Likelihood:** the probability the trained model predicts the target to be present at a particular location in a frame. This value reflects the confidence of the predictions and ranges between 0 and 1, with 1 indicating high confidence and 0 indicating low confidence.

**Generalizability:** the ability of a trained model to accurately predict the location of the target in unseen frames. Good generalizability indicates that the model performs well in unseen frames; poor generalizability indicates that a model performing well in training does not perform well in unseen frames. Poor generalizability is usually related to overfitting (see Test-train loss).

**Test-train loss:** the difference between test RMSE and train RMSE (test RMSE − train RMSE). A small test-train loss indicates the model generalizes well to unseen data. A large test-train loss suggests overfitting, where the model has learned features too specific to the training frame, rather than general visual features such that it only performs well in training frames but not in novel frames.

**(4) Applying the trained model to new videos**

**Inference/predictions:** applying the trained model to new videos to predict x-and y-coordinates and likelihoods for each labeled body part.

**CSV files:** plain-text, comma-separated tables DLC outputs. This file contains per-frame x, y coordinates and likelihoods for easy import into analysis tools.

**HDF5 files (H5):** binary, hierarchical data format DLC uses for efficient storage of large pose-estimation datasets with metadata.

#### BOX 2: User Input Parameters for DLC

DeepLabCut offers a range of user-adjustable input parameters that control project setup, model training, and post-training analysis. These parameters are set at two stages: (1) the project configuration file (config.yaml), which is created automatically when a new project is initialized, and (2) as arguments passed directly to DLC function calls in Python or through the DLC graphical user interface. Below, we describe the parameters most relevant to users beginning a new project. For a complete list of parameters, see Nath et al. (2019).

(1) **Input parameters specified in config.yaml file**

**start/stop:** fraction of the video at which extraction of frames starts and stops. For the entire video start=0, stop=1.

**numframes2pick:** how many frames per video to extract for labeling.

**skeleton:** defines connections between bodyparts for visualization. No effect on training or performance.

**p-cutoff:** likelihood threshold for filtering predictions during analysis (default= 0.1). It is used in post-training analysis and does not affect training. Predictions below this threshold are marked as unreliable. Lower values (e.g., 0.1) retain more predictions, including uncertain ones. Higher values (e.g., 0.9) retain only confident predictions but may discard valid data.

**dotsize**: size of the annotation dot during manual labeling. This affects labeling accuracy and has no effect on training. Large dotsize limits the precision with which ground-truth keypoints can be placed on small objects. **alphavalue:** a parameter for visualization of model predictions only. It controls the transparency of label dots and prediction markers in DLC’s evaluation plots and labeled frames. A value of 0 is fully transparent and 1 is fully opaque.

**colormap:** color scheme for visualizing labels and predictions. No effect on training or performance.

**Iteration:** a number tracking which refinement cycle the project is on. A value of 0 indicates first round of labeling, training and evaluation. A value of 1 indicates a new model is being trained using additional or refined labels based on evaluation of predictions in iteration 0. In general, higher values signify further refinement. It determines which model folder DLC reads and writes to. Note that the term iteration in a config.yaml file is different from the term training iteration (mentioned in Box 1B; training iteration can be adjusted as *maxiters* in).

**net_type:** the backbone network architecture used for training. Options include resnet_50, resnet_101, resnet_152, mobilenet_v2_1.0, mobilenet_v2_0.35, efficientnet-b0 through b6. For resnet_n, the number n indicates the number of layers in the backbone of the network. The choice can affect training time per iteration, GPU memory requirement, inference speed when analyzing new videos, and prediction accuracy. For example, a deeper backbone almost always requires higher GPU memory and training time. It can improve prediction accuracy to a certain point, but too many layers may result in diminished return or even overfitting of the model. Generally, deeper backbone networks are recommended for more complex tasks.

**TrainingFraction:** fraction of labeled frames to be used for training (referred to as Training proportion in this paper). Higher fractions allocate more labeled frames for training and fewer for testing.

**default_augmenter:** specifies the pipeline used for data augmentation (the process of creating modified versions of training frames to increase diversity of the examples available for the neural network) during training. Options include ‘default’ (scaling and cropping only), ‘imgaug’ (rotation, scaling, motion blur; recommended for most projects), and ‘tensorpack’ (similar to imgaug but with multi-CPU support). More diverse augmentation can improve model generalizability, particularly for small training sets.

**snapshotindex:** controls which saved snapshot DLC uses for evaluation and video analysis. A value of 0 uses the first snapshot,-1 uses the last snapshot, and “all” uses all the snapshots.

**Batch_size:** number of frames processed simultaneously per training step (default=1) or per analysis step (default=4). Larger values speed up processing but require more GPU memory.

*(continued on next page…)*

**cropping:** the coordinates (x1, x2, y1, y2) used to crop video frames.

**Shuffle:** a randomized split of labeled frames into train and test data. It is recommended to run training on multiple shuffles to ensure that model results are not dependent on a single split.

(2) **Additional input parameters set through the DLC interface or as arguments in the training function call maxiters:** total number of training iterations. It affects prediction accuracy, training time, and GPU usage duration. Beyond the convergence plateau, additional iterations would likely result in diminished return in model performance. **saveiters:** how often model snapshots are saved (e.g., every 100,000 iterations). This affects disk space use (each snapshot is ∼100 MB depending on the backbone) and the granularity of your evaluation. Lower values save snapshots more often, providing finer grained evaluation, but consume more disk space.

**Displayiters**: how many cross-entropy loss values are printed to the console or log. This value has no effect on training. A higher value offers finer grained resolution in the cross-entropy loss curve.

## Methods

### Video acquisition and processing

Sexually mature, 7-10 days post final molt females from a lab colony of the Hawaiian cricket *Laupala cerasina* were used to obtain videos. Males of *Laupala cerasina* produce species-specific long-range songs to attract females and females orient and walk towards males that sing her preferred song (Shaw and Herlihy, 2000). Behavioral trials were conducted in a cylindrical arena (1 m diameter, 60 cm tall) inside a sound isolation chamber (ETS-Lindgren, Cedar Park, USA) maintained at 20°C. The wall of the arena was covered with black felt and the bottom of the arena was lined with a white plastic interior wall panel (Parkland Performance, Middlebury, USA) (Fig. 2). A speaker (Gallo Acoustics, Scotland, UK) was placed behind the arena wall to broadcast the stimulus song. The speaker was calibrated to 80 dB, 0.5 m from the speaker using a sound level meter (Type 2250, Bruel & Kjaer, Nærum, Denmark) before trials each day. Because *Laupala* crickets are active during the day, the sound chamber was lit by overhead LED lights during the behavioral trials.

**Figure 2.**
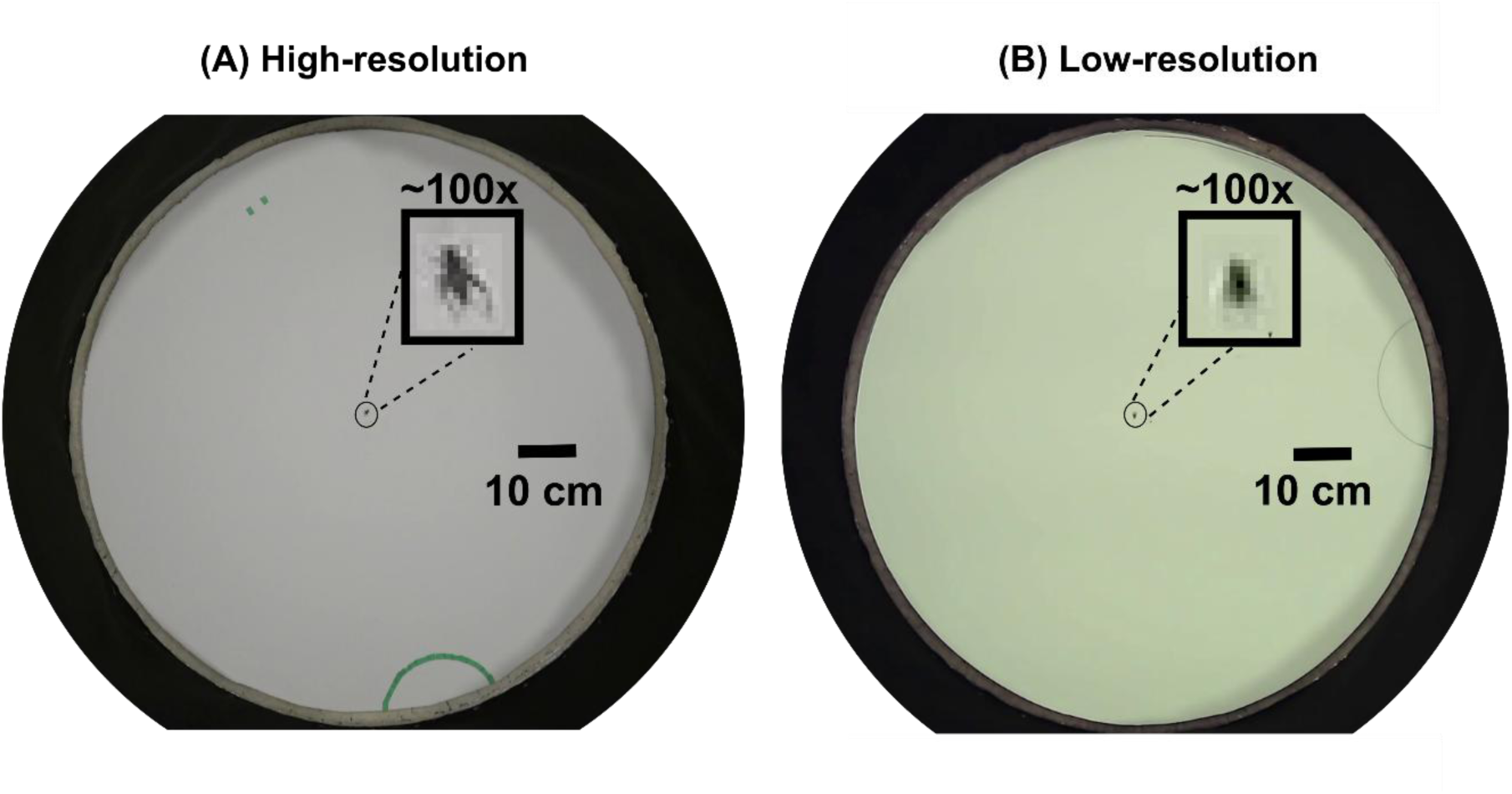
Representative video frames from (A) high-resolution (1920 × 1080 pixels) and (B) low-resolution (1536 × 960 pixels) top-view recordings of behavioral trials in an arena of 1 m diameter. Insets show focal crickets (∼1 cm body length) magnified 100 × at native pixel resolution, spanning approximately 10 and 7 pixels in high-and low-resolution frames, respectively. The black boundary represents the vertical arena wall covered with a black felt cloth. The semicircles demarcate the 10-cm radius response zone around the active speaker. Scale bars = 10 cm.

Each trial consisted of a 1-min habituation period and a 5-min testing period. During the habituation period, a single female was placed under a plastic cup in the center of the arena with the stimulus song playing. After habituation, the plastic cup was lifted through a fishing wire from outside the chamber to release the female, commencing the testing period. Movements of the focal female were recorded using an overhead camera mounted on the ceiling of the chamber, approximately 2 m from the floor of the arena. High-resolution recordings were made using a Sony FDR-AX53 camera with an Exmor R CMOS sensor, a ZEISS Vario-Sonnar T* 20× optical zoom lens (Sony Corporation, Tokyo, Japan) at 30 frames per second (fps). Low resolution videos were made with a ELP USB camera with an Aptina MI5100 CMOS sensor (Shenzhen Ailipu Technology Co., Ltd., Shenzhen, China) 2.8–12 mm varifocal lens at 30 fps. We recorded twenty trials using eighteen females under each resolution, resulting in a total of 40 videos for tracking.

All videos were processed before they were used for movement tracking. We first cropped areas outside the arena while maintaining the aspect ratio to improve processing time. After cropping, the dimensions for the high-and low-resolution frames were 1920 ×1080 and 1536 ×960 pixels, respectively (Fig. 2). Video frames representing scenarios where the focal female was not visible in the frame, such as when the female was underneath the plastic cup and when rare connection glitch produced a black screen in high resolution recordings, were excluded from analyses.

### Software and hardware setup

We used DeepLabCut version 2.3.3 and a Resnet-50 based model (see net_type in Box 2; Mathis et al., 2018; Nath et al., 2019) installed on a Dell OptiPlex desktop computer with a NVIDIA GeForce RTX 3090 graphic processing unit (GPU, Box 1B).

### Project setup and manual labeling

For both high and low-resolution recordings, we extracted and manually labeled the focal animal in 10 frames per video (see *numframes2pick* in Box 2). Frame extraction was done using K-means clustering (Box 1B) to maximize diversity of frames within the training sets. After excluding scenarios where crickets were invisible, there were 195 and 197 manually labelled frames for high-and low-resolution recordings, respectively. For high-resolution frames, the experimenter was able to visually distinguish between the head and the abdomen of the cricket. Hence, two distinct labels were manually placed in the center of the head and abdomen and labeled as “head” and “body”, respectively (see *bodyparts* in Box 2). For low resolution frames, body parts were not distinguishable visually and hence, a single “body” label was placed in the center of the cricket. For a full list of the user defined parameter values see Box 2.

### Training design and parameter setting

We focused on examining the effect of two input parameters on performance of the program: training proportion (% of manually labelled frames used for training; Box 1B) and the number of training iterations (Box 1B). Higher training proportion and iterations typically correspond to low error rates (i.e., better performance) but also involve higher time/GPU demand. At each recording resolution, we constructed models with three training proportions (95%, 80% and 50%). For each model, we saved snapshots at six training iterations (100,000, 200,000, 300,000, 400,000, 500,000, and 600,000). This resulted in 18 combinations of the two parameter values for each resolution.

### Model evaluation

To investigate the effect of recording resolution and input parameters on model performance, we evaluated performance using four evaluation parameters: training convergence, prediction error, prediction certainty, and generalizability. All evaluation parameter values were taken from the test frames unless noted otherwise and analyses of these parameters were performed in R v4.5.3 (R Core Team, 2026). Additionally, we recorded the timestamps of snapshot files to calculate training time for each of the 18 model configurations across both recording resolutions. The training time was used as a measure of computation cost to achieve a certain level of performance.

### Training convergence

We fitted a three-parameter exponential decay model (loss = *a* · exp (−*b* · x) + *c*) using nonlinear least squares (“minpack.lm” package in R; Elzhov et al., 2016) with the number of training iterations as the independent variable and cross-entropy loss as the dependent variable. The parameter *c* represents the asymptotic plateau of the fitted curve. A 95% confidence interval for *c* was computed by bootstrapping the cross-entropy data 1,000 times (seed = 42; “boot” package in R; Canty & Ripley, 2017). The training convergence point was defined as the lowest iteration at which the fitted curve entered the 95% confidence interval of the asymptotic plateau parameter. Because model snapshots (Box 1B) were saved at 100,000-iteration intervals (see *saveiters* in Box 2), the convergence point was rounded up to the nearest 100,000 iterations for which data for evaluation were available.

### Prediction error

Model performance was assessed along two dimensions: prediction error and prediction certainty.

To evaluate prediction error, we calculated the root mean square error (RMSE) in pixels between the model’s predictions and the manually labeled ground-truth keypoints for both test (test RMSE) and train frames (train RMSE). RMSE values were calculated in two ways: with all predictions included (no p-cutoff) and with predictions having likelihood below 0.1 excluded (p-cutoff applied; see Box 2). For high-resolution videos, we calculated test and train RMSE for both head and body labels; RMSE values were similar for both labels (see Supplementary Materials) and therefore, only body labels were used for comparison with low-resolution frames. RMSE was calculated using the formula: 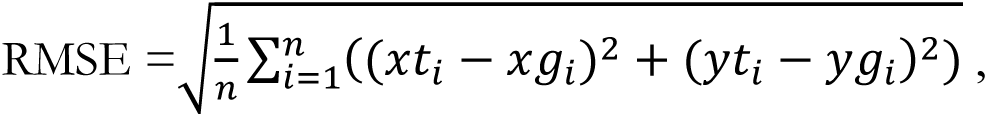 where *xt_i_*, *yt_i_* and x*g_i_*, y*g_i_* are the x and y coordinates for the model predictions (t) and the ground-truth (g) keypoints, respectively for the *i*th test frame, and *n* is the total number of frames. DLC typically provides a csv file with both test and train RMSE (with p-cutoff applied and with no p-cutoff) during the model evaluation step (Fig. 1D and Box 1B) which was used for further analyses.

Our evaluation metric primarily focused on test RMSE with p-cutoff applied (i.e., excluding predictions with extreme low confidence) as prediction error for test frames provides a robust estimate of performance on novel data. Because RMSE in the unit of pixels are specific to the recording resolution and size of the focal animal, making it difficult to interpret and compare across different recordings, we also calculated relative RMSE, which is the pixel RMSE expressed in the unit of the average body length of the subject (relative RMSE = test RMSE / average body length). Body length was measured as the distance from the anterior tip of the head to the posterior tip of the abdomen (cerci not included) and body area was measured as the area of the polygon delineating the outline of the cricket, both using ImageJ v1.54g (Schneider et al., 2012) at 1:1 zoom from one randomly selected frame per video for each recording resolution. To provide a baseline to compare model RMSE to, we calculated a within-human RMSE from two sets of manual labels in the same frames labeled by the same experimenter.

### Prediction certainty

Certainty of model predictions was assessed by the proportion of test frames with likelihood (Box 1B) ≥ 0.9 for model.

### Generalizability

We assessed generalizability of models using test-train loss (Box 1B), calculated as test RMSE minus train RMSE after applying the p-cutoff. A small test-train loss, regardless of the sign, indicates that the model generalizes well to novel frames and a large positive test-train loss suggests overfitting (i.e., the model has learned features specific to the training frames that do not transfer to novel frames well, rather than general visual features). Similar to the prediction error, we also provide a biologically interpretable measure of relative test-train loss measured in the unit of average body length.

### Optimal model selection

To compare performance between resolutions and diagnose tracking errors, we selected one optimal model for each resolution. Our selection criteria included two aspects that are in a trade-off: model performance and computational time. We selected model configurations with acceptable performance using the following criteria: (1) training iterations at or beyond the convergence point; (2) relative test RMSE ≤ 0.33 (i.e., test RMSE within one-third of average body length); (3) proportion of test frames with likelihood at or above 0.9 ≥ 95%; and (4) the absolute value of relative test-train loss ≤ 0.33. The optimal model for each recording resolution was the one with the shortest training time among models meeting all four criteria (Fig. 1E). This cost-conscious approach considers the trade-off between performance and computational cost and prioritizes computational efficiency among models with adequate tracking performance, rather than selecting the single best-performing model regardless of cost.

### Data analysis using the optimal model

We applied the optimal model for each recording resolution to all 20 videos and made inference using the optimal model. We assessed and compared likelihood for the two recording resolutions. Note that for inference, because human labeled ground truth was not available, no assessment about prediction error can be made. In addition, we inspected frames with likelihood values below 0.9 to characterize scenarios associated with reduced prediction confidence. We derive categories of errors based on recurring visual patterns observed across multiple videos and assigned each low-likelihood frame a category.

## Results

On average, the cricket body length was 10.13 ± 0.11 (mean ± SE) pixels with an average area of 71.7 ± 2.45 px² (n = 20) in high-resolution frames and 7.03 ± 0.18 pixels with an area of 36.6 ± 1.91px² (n = 20) in low-resolution frames. Across frames, the cricket occupied approximately 0.009% and 0.01% of the area of the bottom of the arena for high-and low-resolution recordings, respectively (Fig. 2).

## Model evaluation

### Training convergence and training time

Cross-entropy loss followed an exponential decay pattern across all training proportions for both resolutions (Fig. 3). For high-resolution recordings, convergence occurred progressively earlier as training proportion increased, with cross-entropy loss plateauing at 171,822, 132,843, and 131,662 iterations for 50%, 80%, and 95% training proportions, respectively (Fig. 3A-C). For low-resolution recordings, there was no consistent relationship between training proportion and convergence timing, with cross-entropy loss plateauing at 93,273, 106,266, and 90,320 iterations for 50%, 80%, and 95% training proportions, respectively (Fig. 3D-F). Based on these values, the convergence threshold for high-and low-resolution recordings was set at 200,000 and 100,000 iterations, respectively. Each 100,000-iteration training segment required approximately 3 hours for high-resolution and 2 hours for low-resolution models.

**Figure 3.**
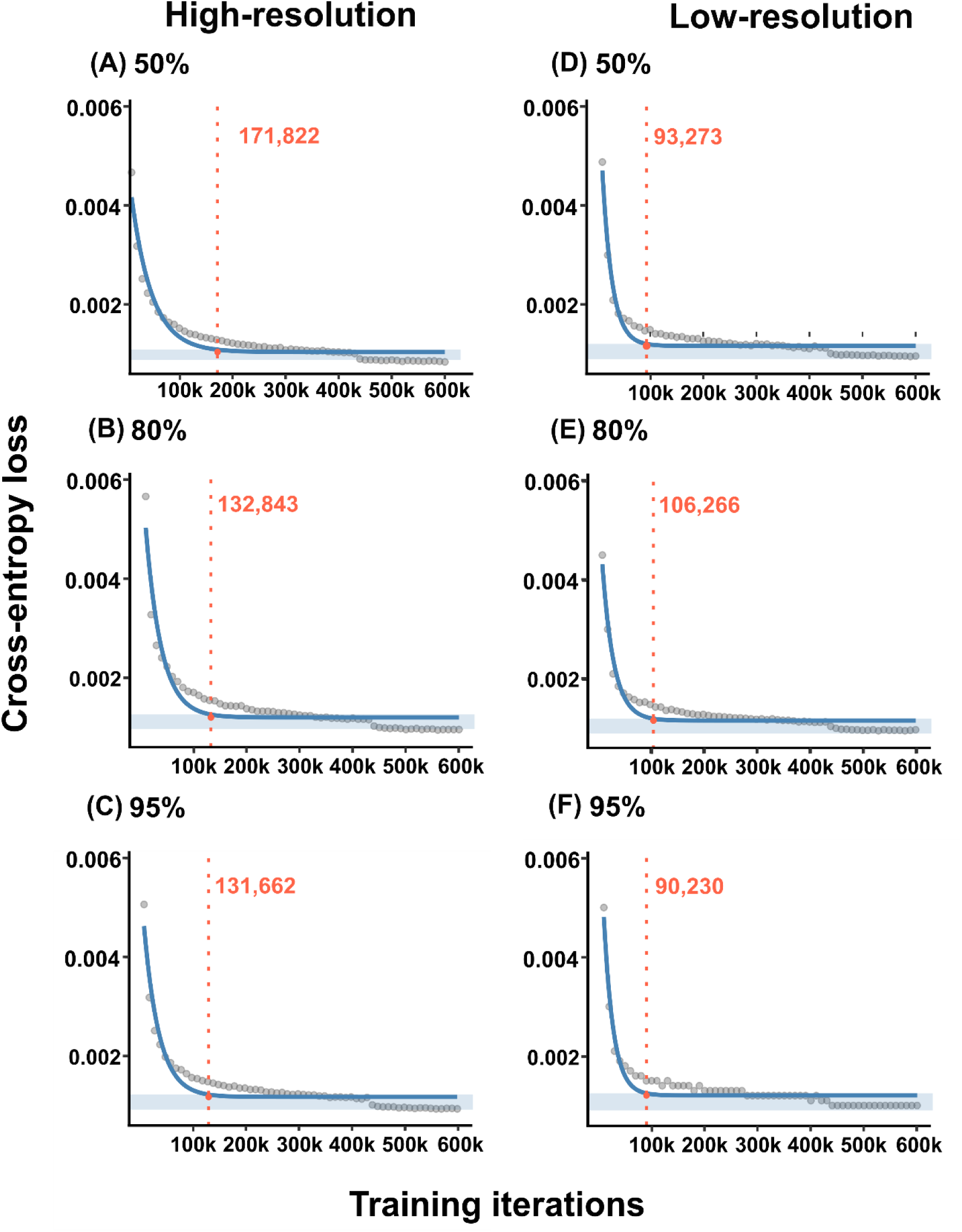
Cross-entropy loss across training iterations for high-resolution (panels A-C) and low-resolution frames (panels D-F). The gray points are raw cross-entropy loss data and solid blue lines are fitted exponential decay curves. The light blue shaded areas correspond to 95% confidence intervals for the plateau estimates of the fitted curves. Red dashed lines indicate the lowest iteration number at which the cross-entropy loss reaches the 95% confidence interval.

### Prediction error

For high-resolution recordings, the test RMSE for post-convergence models ranged between 1.61 and 2.44 pixels, translating to relative RMSE between 0.16 and 0.24 of the average body length of the cricket (Fig. 4A-C; Table 1), all below the 0.33 acceptance threshold. Relative RMSE values did not change when a p-cutoff of 0.1 was applied (Table 1). There was no consistent pattern of change in relative RMSE with training iterations or training proportion (Table 1).

**Figure 4.**
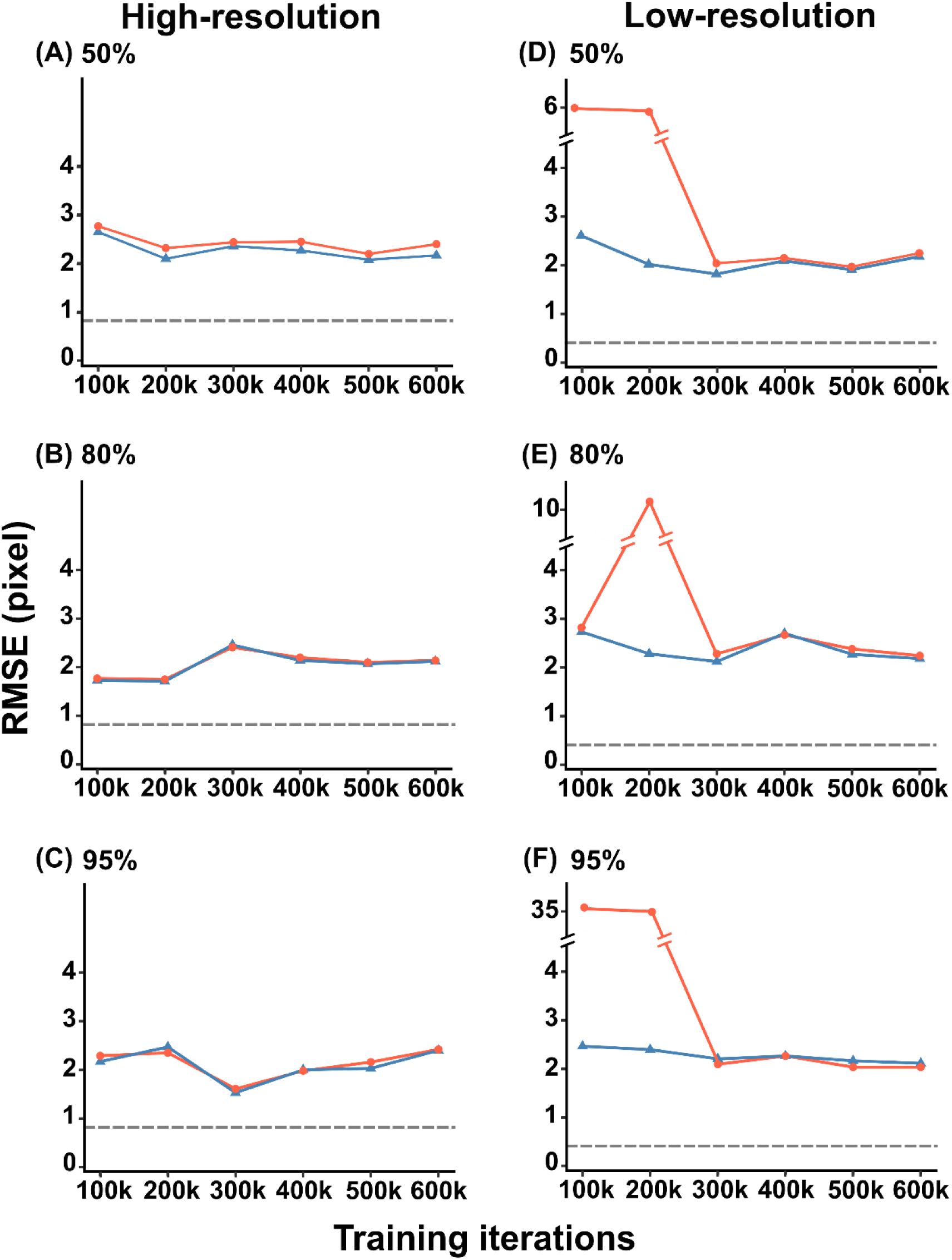
Root Mean Square Error (RMSE, Box 1B) of predicted locations of the focal cricket in test (orange) and train (blue) frames at six iteration numbers for high-resolution (panels A-C) and low-resolution frames (panels D-F). Values are shown after a p-cutoff of 0.1 is applied. The grey dashed horizontal lines represent within-human RMSE for the given recording resolution. Note that for low-resolution models, high RMSE values in early iterations are shown on discontinuous Y-axes, marked by break symbols on the Y-axis and lines connecting data points.

**Table 1.**
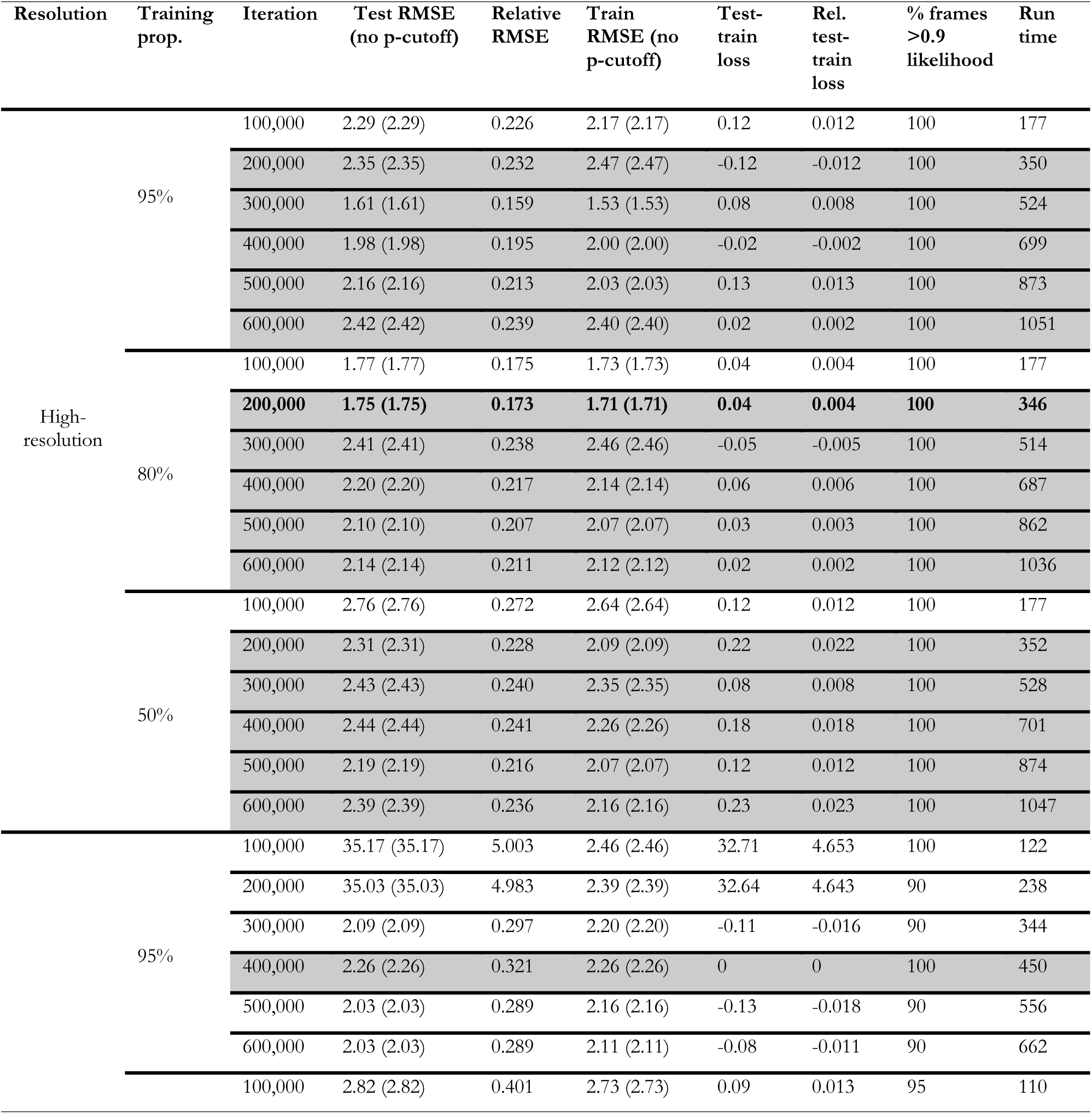
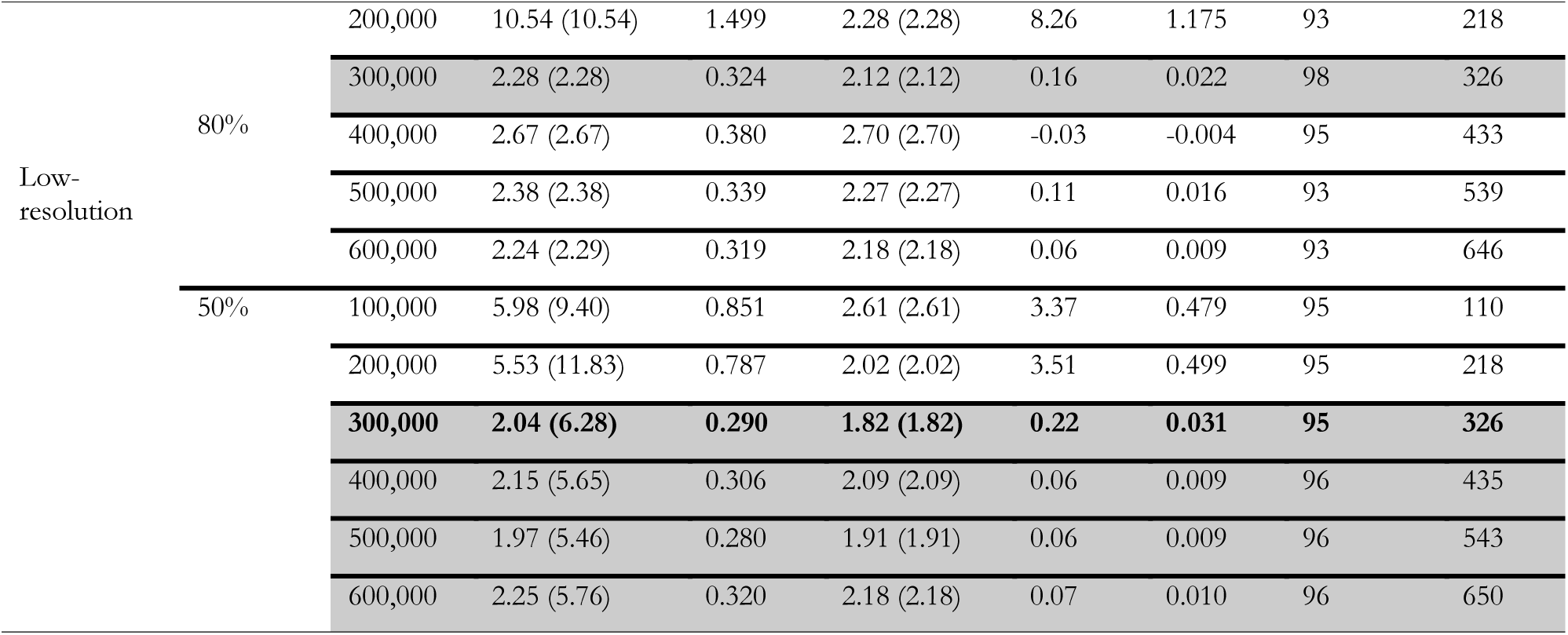
Model performance and generalizability across all training configurations. Grey shaded rows represent model configurations that met the convergence, prediction error, prediction certainty and generalizability criteria. Bold values appear in rows representing the selected optimal model configuration.

For low-resolution recordings, test RMSE for post-convergence models ranged between 1.97 and 35.17 pixels, translating to relative RMSE values ranged from 0.28 to 5.00 of the average body length of the cricket (Fig. 4D-F; Table 1). The anomalously high values of test RMSE corresponded to the 95% training proportion at 100,000 and 200,000 iterations (Table 1; Fig. 4D). Unlike the high-resolution recordings, relative RMSE values decreased when a p-cutoff was applied for multiple parameter combinations, indicating co-occurrence of frames with low-likelihood and high prediction error (Table 1). Relative RMSE generally decreased before 200,000 iterations and there was no consistent pattern of change in RMSE with training iterations or training proportion beyond 200,000 iterations, with relative RMSE ranging between 0.28 and 0.38 of the average body length of the cricket (test RMSE: 1.97–2.67 pixels). All but two values in this range fell within the 0.33 acceptance threshold.

The within-human RMSE was 0.82 pixels (0.08 of the average cricket body length) for high-resolution and 0.41 pixels (0.06 of the average cricket body length) for low-resolution frames. Model errors were consistently higher than within-human RMSE for all models under both resolutions.

### Prediction certainty

All high-resolution models had more than 95% of test frames with likelihood ≥ 0.9 (Table 1). For low-resolution recordings, the number of models meeting the 95% prediction certainty criterion varied with training proportion: all 6 models at 50% training proportion met the criterion, compared to 3 of 6 at 80% and 2 of 6 at 95% training proportion. However, among models that met the criterion, the proportion of high-likelihood frames showed no clear pattern of change with either training proportion or iteration count.

### Model generalizability

For high-resolution recordings, the absolute value of relative test-train loss (test-train loss / average body length) was within 0.33 for all post-convergence models (Table 1). For low-resolution recordings, relative test-train loss overall decreased beyond 200,000 iterations (except for 80% training proportion at 100,000 iterations). Beyond 200,000 iterations there was no obvious pattern of change in relative test-train loss with training proportion or iterations.

### Optimal model selection

For high resolution recordings, all but three model configurations met all four performance criteria (Table 1, shaded rows). Among these, the model configuration with 80% training proportion and 200,000 iterations had the shortest training time and was chosen as the optimal model for analyzing all high-resolution videos. Estimating from the test frame set, this configuration had a relative RMSE of 0.17 of the average cricket body length, relative test-train loss of 0.00, and 100% of frames with likelihood ≥ 0.9, with a training time of 346 min. For low resolution recordings, six out of 18 model configurations met all performance criteria (Table 1, shaded rows). Among these, two had the same least run time (50% training proportion and 300,000 iterations and 80% training proportion and 300,000 iterations). We chose 50% training proportion and 300,000 iterations as the optimal model as it required less manual labelling time (∼ 100 frames for 50% versus ∼160 for 80%). This model configuration achieved relative RMSE of 0.29 of the average cricket body length, relative test-train loss of 0.031, and 95% of frames with likelihood ≥ 0.9, with a run time of 326 min.

## Data analysis using the optimal model

A total of 139,781 high-resolution frames across 20 videos were analyzed by the optimal model (example traces in Fig. 5A). Total running time was roughly 90 min, translating to on average 69 s per 1 min of high-resolution video recorded at 30 fps. 98.68% of these frames had likelihood ≥ 0.9. Frames with likelihood < 0.9 (red dots in Fig. 5A) were categorized into one of the four categories (Fig. 6): cricket positioned within shadow of the arena wall (60.5%), cricket on top of the marking of the response zone (13.9%), cricket on the black felt wall of the arena (14.1%), and no visually discernible cause (11.5%).

**Figure 5.**
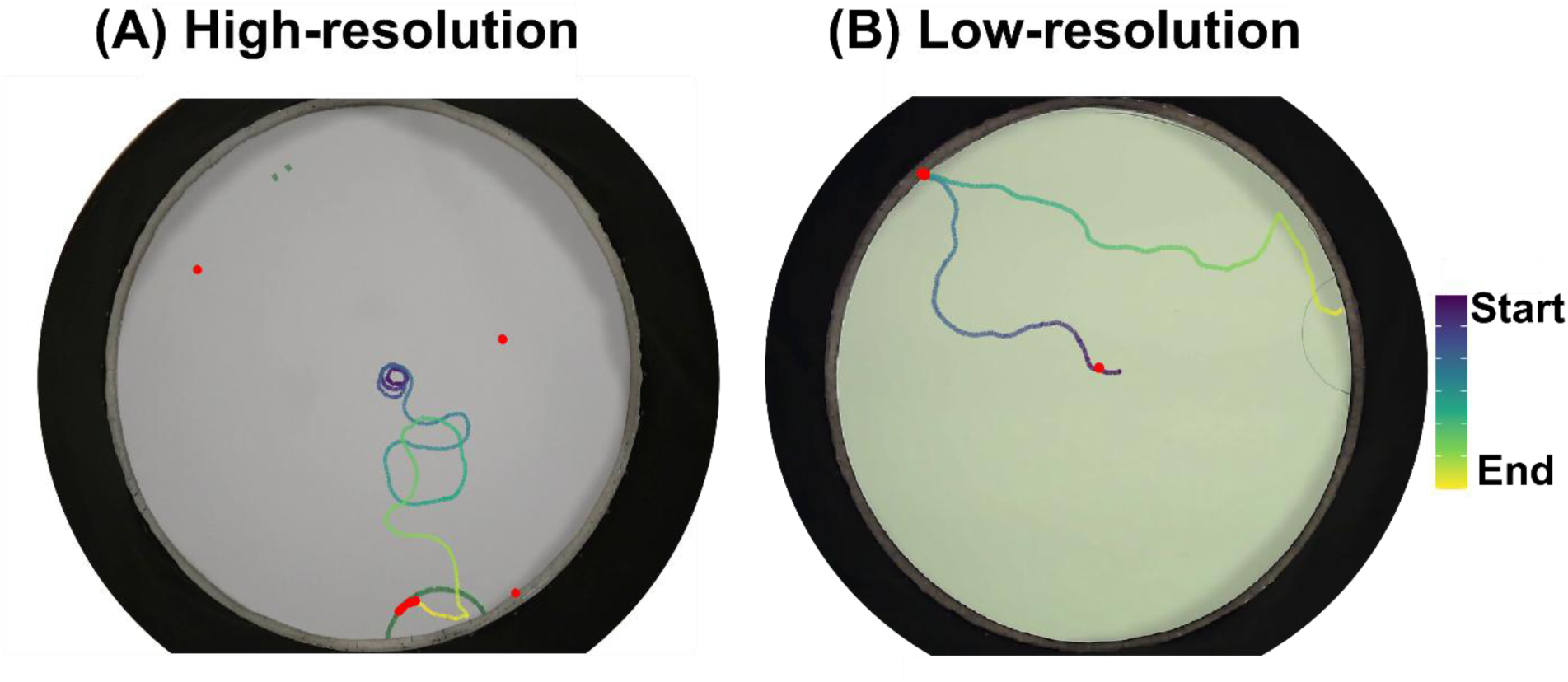
Representative movement traces from automated DLC tracking for (A) high-resolution and (B) low-resolution recordings. Color gradient indicates time progression (dark purple = start of trial, yellow = end of trial). Red dots indicate frames with likelihood < 0.9. The three red dots in (A) displaced from the continuous movement path indicate erroneous model predictions.

**Figure 6.**
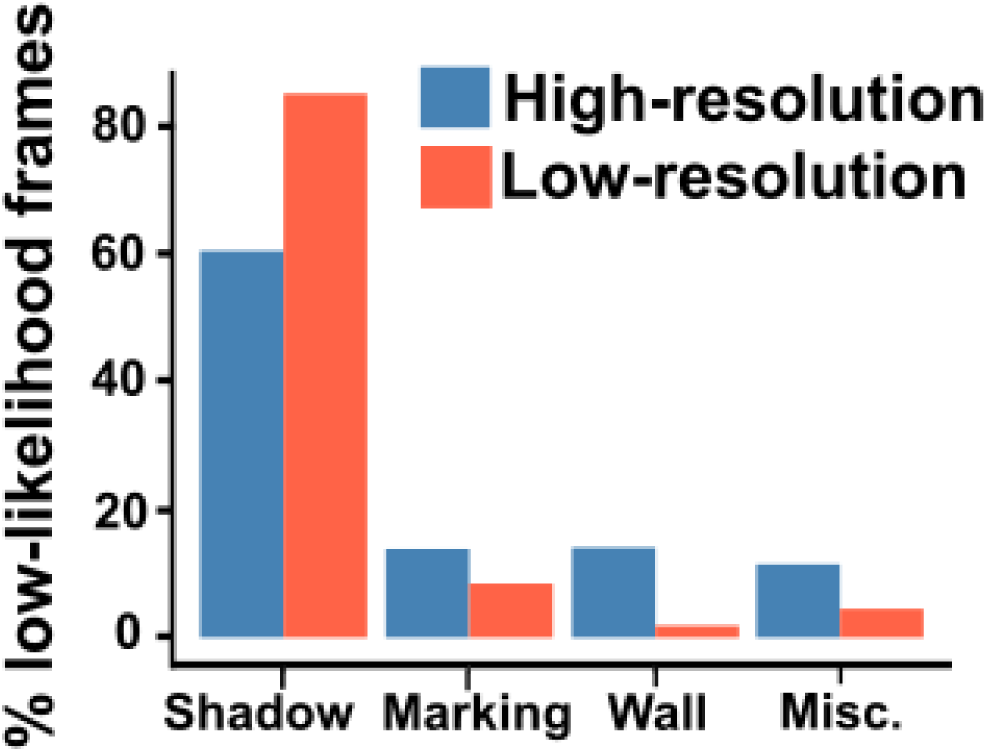
Categories of frames with low likelihood values from predictions in 20 high-and 20 low-resolution recordings made by the corresponding optimal model. Shadow, Marking and Wall correspond to scenarios where the focal cricket was positioned within the shadow cast by the wall of the arena, darker-colored markings on the arena floor to demarcate the response zone, and on the dark arena wall, respectively. Uncategorized correspond to scenarios with no obvious visual discernable cause for low likelihood values.

For low-resolution recordings, 227,991 frames across 20 videos were analyzed using the optimal model with a run time of 95 min (Fig. 5B shows an example trace). This translates to ∼45 s per 1 min of video recorded at 30 fps. 96.61% of frames had likelihood ≥ 0.9. Of the low-likelihood frames (red dots in Fig. 5B), in 85.1% of them the focal cricket was within the shadow cast by the wall of the arena, in 8.5% the cricket was on top of response zone markings, in 1.9% the cricket was on the black felt wall, and in 4.5% there was no visually discernible cause.

## Discussion

Machine learning-based methods have dramatically increased the analytical throughput of behavioral tracking in a variety of fields. Despite the advantages, implementation of ML-based tracking tools is met by several barriers, including a steep initial learning curve for the users and lack of validation data in diverse contexts. This paper extends the contributions of other comprehensive guides to DLC (Nath et al., 2019) by providing a laymen primer for workflow and vocabulary of ML-based tracking. Additionally, we evaluate DLC’s performance in a common but unvalidated scenario where the focal subject is relatively small. We also compare model performance between high and low recording resolutions, corresponding to high and low budget in the recording devices and offer suggestions about choosing user-defined input parameter values for both resolutions.

Our data showed that DLC can achieve adequate performance when the target size is relatively small. In this study, the focal animal occupied < 0.01% of the arena area under both resolutions, ratios substantially smaller than those reported from any previous evaluation of DLC. The optimal models for both high-and low-resolutions achieved test RMSE less than one-third of the average cricket body length (Table 1). Many animal behaviors like mate attraction, predation, foraging, parenting, and schooling are often quantified using gross spatial patterns like orientation, distance traveled, speed, presence/absence in a pre-designated area, and positive/negative attraction to a stimulus, and for such metrics, tracking errors less than one-third of the body length are unlikely to be consequential for either accuracy or precision of tracking. The optimal models also achieved low errors with high certainty in their predictions. The optimal models yielded 100% and 95% of test frames with a likelihood ≥ 0.9 for high-and low-resolution recordings, respectively.

Additionally, the optimal models exhibited high generalizability with as the test RMSE and train RMSE remained similar with error differences as small as 0.4% and 3% of the body length for high-and low-resolution recordings, respectively. At the inference stage when the entire dataset was analyzed, the optimal models again yielded approximately 99% and 97% of predictions with a likelihood ≥0.9 for high-and low-resolution recordings, respectively. These results expand on previous validation data (Chanthongdee et al., 2024; Correia et al., 2024; Hayakawa et al., 2024; Lauer et al., 2022; Mathis et al., 2018; Nath et al., 2019; Sturman et al., 2020) and suggest that with proper choice of input parameter values, DLC can offer satisfactory performance when the relative target size is small. Notably, RMSE from DLC remained slightly higher than human RMSE, although the difference was minimal, suggesting that for objects of this pixel size, automated tracking may not reach human-level precision. Whether this gap can be narrowed through alternative network architectures, augmentation strategies, or better recording conditions remains open.

Critically, our data showed DLC achieved sufficient tracking accuracy within a reasonable timeframe. Running on a NVIDIA GeForce RTX 3090 GPU, training converged and optimal model performance was achieved within 300,000 iterations for all models under both recording resolutions, translating to approximately 6-9 hrs (Table 1). Time required to manually label frames can be highly variable. As a frame of reference, labeling 200 training frames cost roughly 1 hr in our study. Together, the entire training workflow amounts to less than 10 hours, arguably much faster than manual analysis. Although time required for training depends on many factors such as the hardware, the choice of the backbone architecture, and the number of targets to track, information about training time improves accessibility of the program, especially for users with no prior experience, yet such information is rarely disclosed and discussed.

Our data revealed how the proportion of labeled frames used for training and number of training iterations interact to affect convergence and prediction error. For high-resolution recordings, convergence occurred progressively earlier as training proportion increased (Fig. 3a-c). For low-resolution recordings, no such pattern was observed; convergence timing was similar across training proportions (Fig. 3d-f). Prediction error and certainty, in contrast, was largely insensitive to training proportion for high-resolution recordings: across all high-resolution model configurations, relative RMSE values were within 0.33 of the average cricket body length and 100% of frames had likelihood ≥ 0.9 (Table 1). For low-resolution recordings, errors at low iterations were substantially higher than those from the corresponding high-resolution recordings (Fig. 5). Notably, even though all low-resolution models converged before or slightly beyond 100,000 iterations, prediction errors remained high to varying levels at 200,000 iterations, suggesting that small-object tracking with lower-resolution devices may require additional training iterations beyond the convergence point.

The patterns observed in our factorial design have practical implications for users selecting training parameters in their own projects. The data suggests that a cost-conscious approach to model selection, rather than defaulting to maximizing model performance, can substantially reduce computational costs in terms of training time with negligible loss in tracking performance. A useful selection approach involves identifying post-convergence models that meet acceptance thresholds for prediction error (relative RMSE ≤ 0.33), prediction certainty (≥ 95% of test frames with likelihood ≥ 0.9), and generalizability (relative test-train loss ≤ 0.33), then selecting qualifying models with the lowest training time. Applying this framework, we found that for both resolutions, multiple post-convergence models performed remarkably similarly, allowing us to select models with lower training proportions and fewer iterations while maintaining tracking accuracy comparable to the best-performing models.

One consideration that emerged from the data concerns test set size. The anomalously high test RMSE (∼35 pixels) at the 95% training proportion for low-resolution recordings was driven by a single frame in which the model prediction was far from the ground truth within a test set of only 10 frames. Similarly, prediction certainty varied with training proportion for low-resolution recordings. All models at 50% training proportion met the 95% likelihood threshold, compared to only 2 of 6 at 95% training proportion, a pattern consistent with the hypothesis that small test datasets may not offer the most accurate performance assessment because they are less robust to outlier frames. Based on these observations, a training proportion of ≤ 80% is advisable when working with small, labeled datasets (≤ 200 frames), as this yields at least 40 test frames for performance evaluation.

In our analysis, the proportion of low likelihood frames did not differ substantially between the two resolutions, although this proportion is slightly lower for low resolution recordings. On the other hand, across both resolutions, erroneous prediction cases can be generally categorized into three scenarios: low light condition and low background contrast. Among these, the dominant cause was the cricket positioned within the shadows cast by the wall of the arena, reducing lighting within which the target was detected (Fig. 6), followed by frames where the cricket was on markings on the background or the dark arena wall, both of which reduces the contrast between the cricket and the background. Only 4.5–11.5% of low-likelihood frames could not be attributed to an identifiable cause (Fig. 5). The consistency of these categories across both resolutions suggested that light condition and background contrast, rather than recording resolution, is the primary determinant of model confidence in our study. Based on these results, we recommend that considerations about light condition and background contrast to be taken prior to experiments and using strategies such as selecting a background that offers maximum contrast and eliminating shadows may offer more perceivable improvements in tracking performance than investing in a higher-resolution camera.

Across all model configurations, we observed two main differences in performance between the two resolutions. First, low resolution models generally have lower proportion of frames that had high prediction certainty (likelihood), but this difference was not substantial (no more than 10%).

Second, several low-resolution models, all at earlier iterations, exhibited poor performance across aspects, showing high RMSE, low likelihood, and a large test-train loss (Table 1, Fig. 4D-F), indicating that these models had low prediction accuracy and certainty and were overfitted. Applying a p-cutoff prior to evaluation significantly improves all evaluation parameters for those models while parameter for other models remained similar (Table 1). Therefore, for low-resolution recordings, we recommend applying the p-cutoff before evaluating model performance. In contrast, no high-resolution models showed such signs of poor performance and applying a p-cut off did not significantly change evaluation of these models (Table 1). Due to these differences, fewer low-resolution models met all our performance criteria than high resolution models.

Despite these general differences, the optimal models for both high-and low-resolution recordings performed remarkably similarly in all aspects of performance as well as training time (Table 1), suggesting that low-cost equipment such as low-resolution webcams could be sufficient for recording videos to be analyzed by DLC. Using a high-resolution recording device can bring improvements in prediction error, but the cost of the devices is often magnitudes higher and running time for the same length of recording would be longer. We encourage users to evaluate whether the gain outweighs the cost based on their experimental requirements and budget. Most importantly, our data demonstrated that markerless behavioral tracking using low-cost recording devices and open-source programs such as DLC, can be implemented in settings where budget is limited, such as teaching laboratories at secondary schools and colleges, small research laboratories, as well as outreach and public engagement programs. This finding offers exciting implementation possibilities and improves accessibility of ML-based markerless tracking.

In conclusion, our data offers validation for using DLC to track movements in settings where the target is relatively small and provides guidelines for parameter choice and model selection. We also demonstrate that background contrast is a primary consideration for reducing estimation error and that recording resolution has minimum impact on model performance. These findings open the door for implementation of DLC in numerous low-budget settings. Our validation data have one limitation as our videos were acquired in a well-lit lab environment which was an appropriate testing conditions for our diurnal study system. We thus call for studies to investigate DLC’s performance in less favorable conditions, such as subjects moving through low light conditions, obstructions in view, and varying backgrounds.

## Use of AI tools

We used Claude (Anthropic) and Gemini (Google) to improve readability of certain manuscript sections and for assistance with R code development, and figure design. All outputs were reviewed, verified, and revised by the authors, who take full responsibility for the content of this work.

## Supplementary material

## Methods

### Comparison of head and body keypoint labels for high-resolution recordings

For high-resolution recordings, two keypoints were labeled on each cricket: the center of the head and the center of the abdomen (body). To determine whether model performance differed between keypoints, we calculated test RMSE and train RMSE separately for head and body labels across all 18 parameter combinations (3 training proportions × 6 iteration counts). We also computed within-human RMSE separately for each keypoint by comparing the two sets of labels produced by the same experimenter.

## Results

Head and body labels produced similar RMSE values across all 18 parameter combinations (Supplementary Table S1). Post-convergence test RMSE for head labels ranged from 1.86 to 3.00 pixels compared to 1.61 to 2.76 pixels for body labels. The difference between test RMSE for head and body labels were all less than 1 pixel and ranged from-0.76 to 0.26 pixels. Because performance was comparable between keypoints and only body labels were available for low-resolution recordings (where body parts were not visually distinguishable), body labels were used for all cross-resolution comparisons reported in the main text.

---

**Supplementary Table S1.**
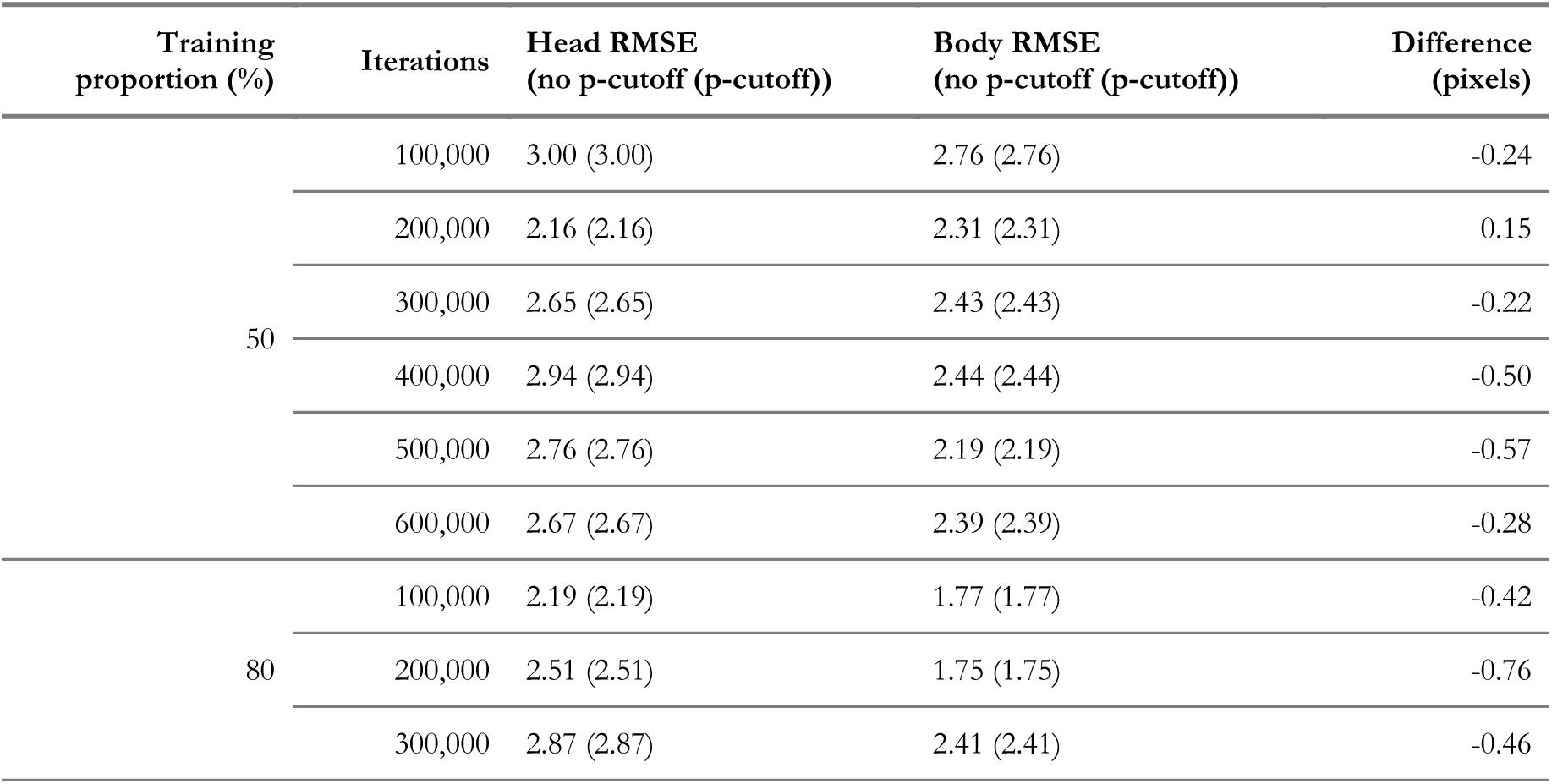
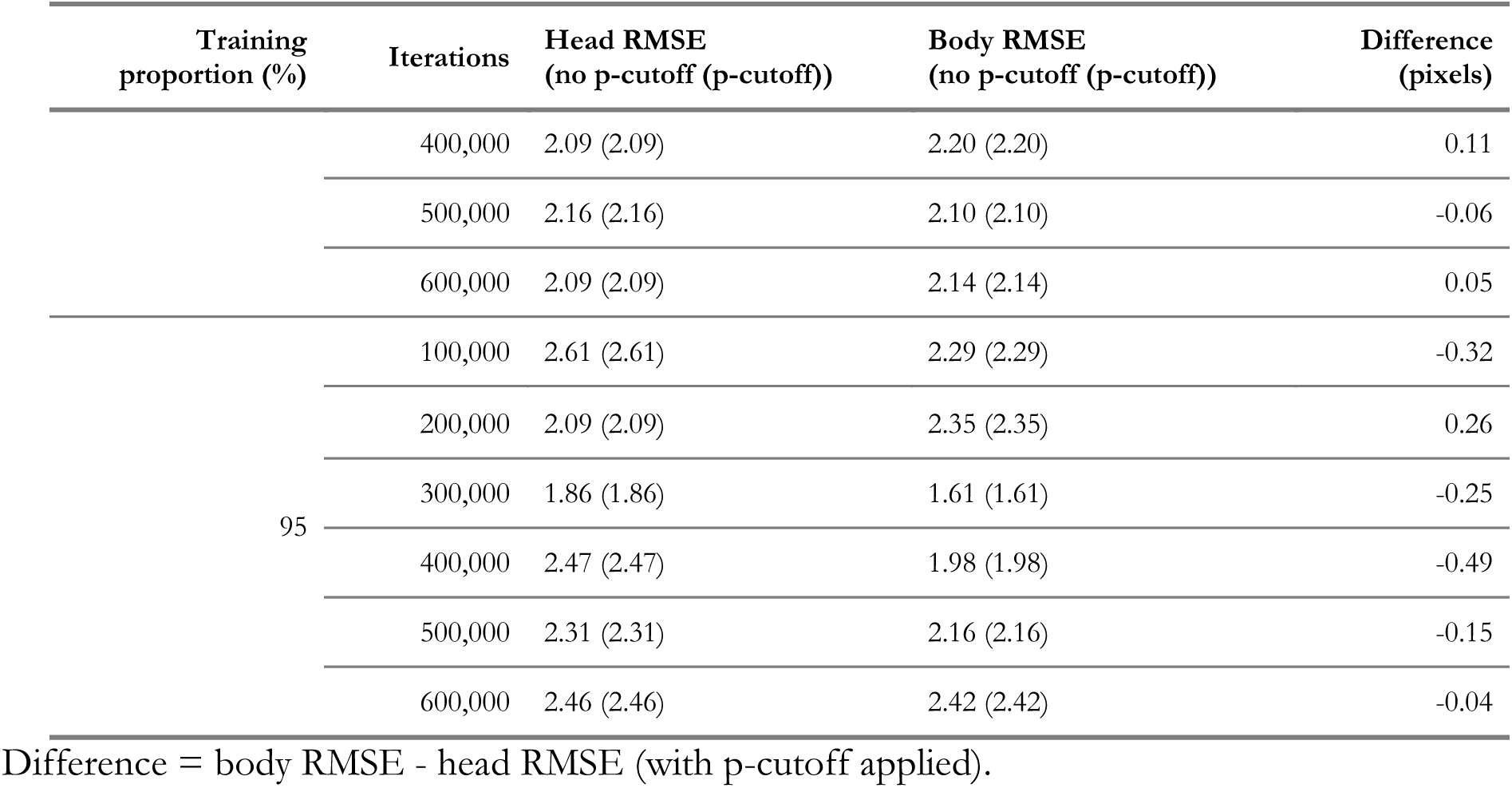
Comparison of test RMSE (pixels) for head and body keypoints across all 18 high-resolution model configurations.

